# Mitigation of radiation damage in biological macromolecules via tunable picosecond pulsed transmission electron microscopy

**DOI:** 10.1101/2020.05.15.099036

**Authors:** Hyeokmin Choe, Ilya Ponomarev, Eric Montgomery, June W. Lau, Yimei Zhu, Yubin Zhao, Ao Liu, Alexei Kanareykin, Chunguang Jing

## Abstract

We report mitigation of electron-beam-induced radiation damage in biological macromolecules using rapid, low-dose transmission electron microscopy (TEM) with a new, tunable, retrofittable picosecond pulser. Damage mitigation strategies historically consisted of sample cryoprotection and ultra-low beam current; ultrafast laser-pulsed systems have shown promise, but with day-long acquisition times. We show the first practical, fast, laser-free tunable system, with acquisition of diffraction series in minutes at 5.2 GHz and 10 pA. This is the largest study to date: two materials (C_36_H_74_ paraffin and purple membrane), two beam energies (200 keV and 300 keV), two independent microscopes (Schottky and LaB_6_), two modes (pulsed and continuous), and unsurpassed repetition rate tunability. Critical dose at room temperature doubled versus continuous beam for ∼100 MHz single-electron repetition rates. Results herald a new class of highly-tunable, ultrafast pulsers with future applications in cryogenic electron microscopy (CryoEM), high resolution single particle imaging, and rapid low-dose TEM.

## Introduction

Transmission electron microscopy (TEM) and its suite of analytical approaches provide unparalleled spatial resolution for characterization of material morphology, structure, and chemistry from sub-nanometer to micrometer scales. But crystalline organic and biological macromolecules are sensitive to damage from the high energy electron beam ^1,2^, typically 60-300 keV. Damage limits achievable resolution and is driven by elastic and inelastic scattering of electron beam by the sample. Elastic scattering of the electron beam by the sample nuclei produces useful diffraction information but can result in atomic displacement or ejection of atoms. Less common inelastic scattering from electron-electron interactions can cause ionization damage (radiolysis). Both scattering processes contribute to electrostatic charging, hydrocarbon contamination and heating of the sample ^3–5^. Radiolysis in particular rapidly damages organic and biological crystal structure; intensity and contrast in diffraction patterns will fade as the same area of the sample is exposed to an increasing dose of beam electrons ^3,6^.

A critical dose can be defined as the threshold at which the maximum intensity of particular reflections of single crystal or powder diffraction drops to 1/*e* of the original value ^7,8^. Critical dose limits the resolution of images, diffraction patterns, and electron tomography ^9^. Among numerous strategies to reduce electron beam damage, cryoprotection ^10–13^ has been highly effective for biological specimens. We discuss a novel and practical implementation of a relatively new damage mitigation technology, ultrafast (picosecond) pulsed beam.

Until recently, critical dose was not thought to depend on how quickly the dose was applied. However, several groups have now reported mitigation of radiation damage via ultrafast electron pulses ^14–17^. These groups used either laser-driven ultrafast electron pulses with kHz to MHz repetition rate, or a pulsed beam of fixed repetition rate produced by GHz radiofrequency (RF) deflecting cavities. Unfortunately, neither approach could achieve highly tunable GHz pulsing to study the impact of high repetition rate picosecond pulse structure on critical dose. Additionally, rigorous comparison of pulsed and continuous beam on one microscope has been challenging on the picosecond scale because the laser-driven cathode modifications replace the normal thermionic or Schottky field emission cathode used for (continuous) operation of the TEM ^18^. The size, divergence, and energy spread of the modified cathode’s pulsed (photoemitted) beam differ from continuous (thermionic or Schottky) emission, a fundamental challenge when comparing pulsed and continuous beam in laser-based pulsed TEM systems. Laser-based systems are further limited to low work function thermionic emitters and do not use higher brightness field emission guns.

We use a laser-free approach to report for the first time rigorous evidence for radiation damage mitigation via tunable, periodically-gated beams in two independently modified stroboscopic TEMs, directly compared to normal continuous beam in the same TEMs. A new technology for pulsed beam generation enables this result; Euclid Techlabs’ STROBO-X00 series allows push-button on/off control of pulsed mode using the original cathode. Both 300 keV thermionic and 200 keV Schottky-emission beam sources show consistent results. Using two model macromolecules, C_36_H_74_ paraffin and purple membrane (bacteriorhodopsin), we calculate a critical dose from a time-intensity diffraction series (fading curves) in selected-area diffraction mode. We show clear improvement in damage mitigation in single-electron-packet mode when an equivalent dose is delivered with a periodic tunable-width picosecond gate regulating single-electron arrival at the specimen. Critical dose for pulsed beam improves by a factor of up to 2 at room temperature compared with the continuous random beam for constant dose rate (electron flux). This factor of 2 exceeds laser-based mitigation efforts recently reported at 200 kHz in paraffin ^14^, and at a remarkable ∼100 MHz average single-electron repetition rate, leading to orders of magnitude faster data acquisition on a highly tunable system.

## Results

### Tunable gating for single-electron arrival or pulsed beam

Fig 1a illustrates a pulsed electron beam generated by the new highly tunable Euclid Techlabs pulser ^19,20^: a STROBO-200 on a JEOL JEM-2010F at Brookhaven National Laboratory (BNL), and a STROBO-300 on a JEOL JEM-3010 at the National Institute of Standards and Technology (NIST) ^21^. Each operates at fixed beam energy: 200 and 300 keV respectively.

**Figure 1.**
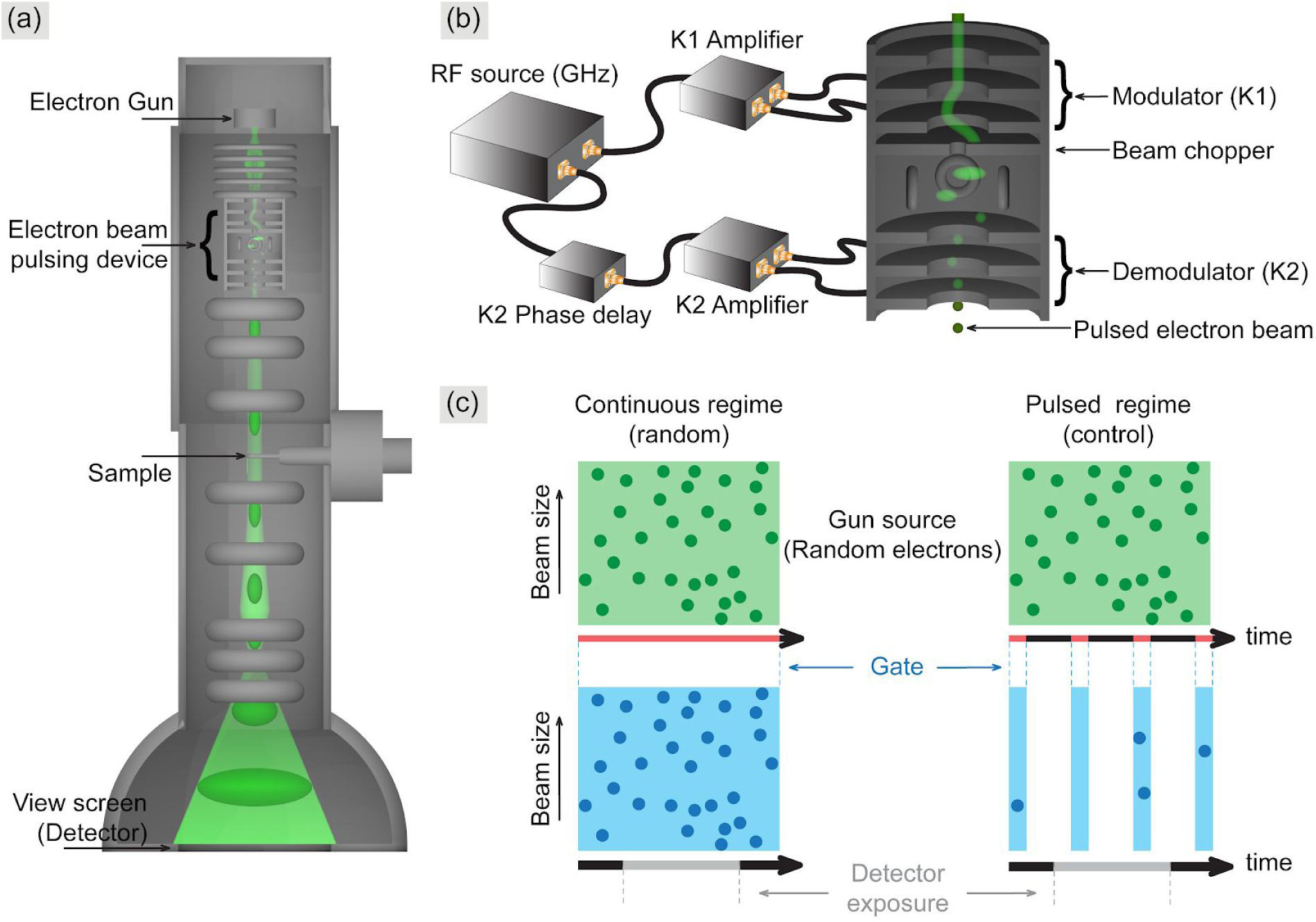
(a) STROBO-X00 series GHz stroboscopic TEM system. (b) 40 MHz - 6 GHz RF drives K1 (and K2 with 180° phase delay) to sweep beam across the chopper aperture, then demodulate transverse momentum. (c) Adapted from ^14^: beam has continuous or pulsed mode; electrons are gated, gate width and frequency are tunable. Detector exposure (time to acquire a single diffraction pattern image) averages across millions of gate periods (not to scale).

Although the gating frequency was fixed in this paper (5.2 GHz), the STROBO-X00 is a broadband device with stroboscopic gating rates from 80 MHz to 12 GHz, since a unique traveling wave architecture overcomes the narrow frequency limitations of GHz resonant cavity deflectors. Gate duration is controlled by frequency, RF power, and chopping aperture diameter.

Direct comparison of critical dose between pulsed and continuous beams using the microscope’s native beam emission process (LaB_6_ filament at NIST, Schottky at BNL) is straightforward, in contrast to laser-based modified cathodes where pulsed beam requires photoemission and continuous beam may not even be an option. Enabling continuous beam (native TEM functionality) is as easy as toggling off RF power to the modulator (K1) and demodulator (K2) (Fig. 1b and 1c). Transverse velocity of the beam is modulated using K1 to produce a transverse RF field sinusoidal in time. The modulated beam, still continuous as it exits K1, is swept across a 0.025 mm diameter chopping aperture which gates the beam. A phase-shifted but otherwise identical RF field in K2 cancels the transverse momentum introduced by K1. Pulsed beam in this work averages tens of pA (single-electron regime): most windows (Fig. 1c) do not have a random electron, but those that do are highly likely to have only a single electron. Each electron in the resulting pulse train is now separated in time from the next electron by at least the gating period. We vary the gate duration in this work (by increasing RF power in K1) while holding the gate repetition rate at 5.2 GHz, and measure average electron arrival rates from 72 − 624 MHz (Table 2). The single-electron packet regime is well-controlled with a short gate width while still achieving high average electron arrival rate (72 MHz at 10% duty cycle) and hence rapid diffraction pattern acquisition. Experiments calibrated in the “continuous random” regime, i.e. normal TEM beam, quantify the critical dose of the beam-sensitive materials in standard experimental conditions. We show these “continuous” baseline results versus “pulsed random” effects by reducing duty cycle (i.e., gate width) and decreasing illuminated area to hold the electron flux constant, for both paraffin and purple membrane, on both the standalone pulsed TEM systems.

### Calibrating radiation damage in organic C_36_H_74_ paraffin crystal with continuous beam

Figure 2a shows a typical diffraction pattern obtained from a C_36_H_74_ paraffin crystal under “continuous random” electron beam using 300 keV TEM, before (top) and after (bottom) exposure to a standard dose of 1000 e^−^·nm^−2^ at room temperature. Diffraction patterns were obtained with 19.39 pA averaged continuous beam current, and the illuminated area was 23.7 µm^2^ so that the electron dose rate on the specimen is 5.09 e^−^·nm^−2^·s^−1^. In Figure 2b, the fading curve data were calculated from the series of diffraction patterns shown in Figure 2a. The grey line is the critical dose (D_cri_), defined as the accumulated electron dose at which the relative intensity (I / I_0_) of the selected diffraction peak decreases to 1/*e* (∼37%).

**Figure 2.**
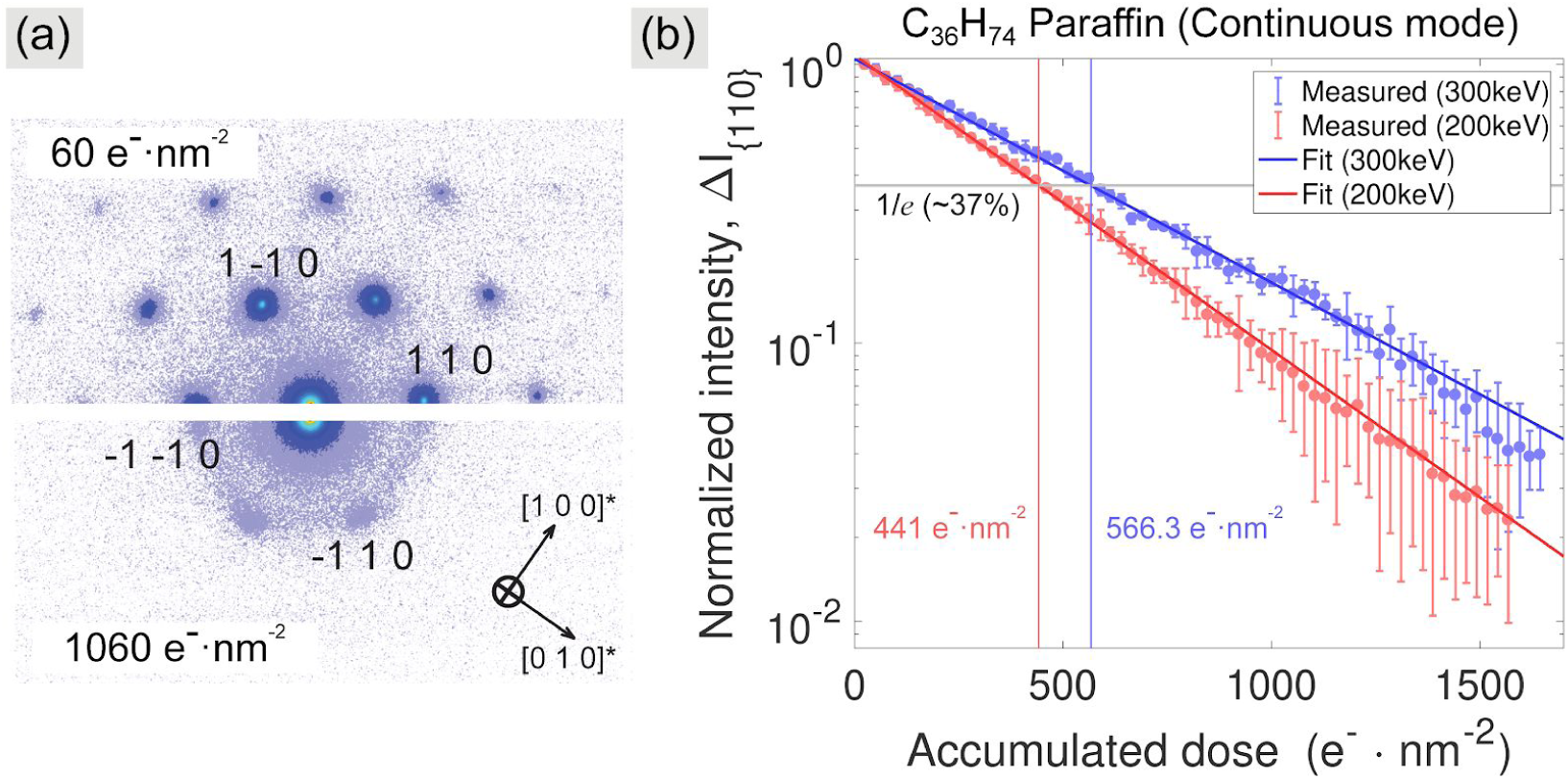
(a) Single crystal electron diffraction on the indexed [001] crystallographic zone-axis for orthorhombic C_36_H_74_ paraffin crystal (a = 7.4 Å, b = 5.0 Å, c = 95.1 Å) ^22^ at room temperature, before and after exposure to 1000 e^−^·nm^−2^ in a 300 keV TEM with a camera length of 300 mm. (b) Fading of the {110} reflections versus total accumulated electron dose are shown for 200 keV (lower, red) and 300 keV (upper, blue), critical dose (D_cri_) in grey. Error bars are standard deviation over 10 measurements on separately prepared samples.

We have fitted each two-dimensional (2D) reciprocal space maps (RSMs) of the {110} reflections to determine the fading curves of the diffraction intensity using a symmetrical 2D Gaussian function. The {110} reflections contain four peaks: (110), (−110), (−1-10) and (1-10). Each RSM involved 8100 intensity values on the mesh of 90 × 90 pixel points. The 2D Gaussian distribution function contains five adjustable parameters: the amplitude of an intensity *A*, the positions of peaks (*x*_0_, *y*_0_) and the peak widths (σ_x_, σ_y_). This means we used 5 model parameters to describe the intensity distribution over 8100 points on each RSM.

The Bragg peak intensity decay as a function of the accumulated total dose is evidence of increasing crystal disorder and irreversible structural damage ^5,7,22^. Dynamic scattering effects are negligible in thin samples. For a given macromolecule and energy we fit the {110} reflections in reciprocal space to 2D gaussians for each image in the dose series, then exponentially fit the fading curve slope for each of the four peaks, and average the result to obtain critical dose. We estimate that the D_cri_ for C_36_H_74_ paraffin in the 300 keV TEM is approximately 540 − 590 e^−^·nm^−2^, as calculated from an exponential fit to the upper curve in Fig. 2b. Likewise, D_cri_ for C_36_H_74_ paraffin in the 200 keV TEM was about 430 − 460 e^−^·nm^−2^, shown in Fig. 2b, lower curve. These “continuous beam” results are comparable to D_cri_ of 340 − 380 e^−^·nm^−2^ reported for C_36_H_74_ paraffin in a 100 keV electron microscope ^13,23,24^, and 440 − 550 e^−^·nm^−2^ for 200 keV ^14^. Further insights from energy dependent damage are deferred to the discussion section; we turn now to the effect of pulsed beam under identical column conditions.

### Mitigation of paraffin radiation damage by tunable pulsed beam

Pulsed beam mode diffraction series showed significant mitigation of C_36_H_74_ paraffin radiation damage. We discuss 300 keV results here and include the same experiment repeated at 200 keV. We first used electron pulses with a 22% duty cycle factor (as defined by the ratio of pulsed to continuous beam current) at 5.2 GHz gate repetition rate. Assuming a flat-top temporal profile, we have 43 ps gate width and 192 ps period. Total accumulated critical dose (D_cri_), flux (dose rate), and all optics except spot size were the same as for the continuous beam (Table 2).

Figure 3a compares the continuous- and pulsed-beam induced radiation damage in paraffin at 300 keV. The diffraction patterns were taken with average electron dose rate 5.11±0.02 e^−^·nm^−2^·s^−1^ (5 second acquisition time). To hold average dose rate (i.e., current density) constant, as we reduced the duty cycle factor from 100 down to 10% (averaged total beam current 100.13 to 10.38 pA), we also reduced the illuminated area proportionally (122.7 to 12.65 µm^2^), all in the single-electron packet regime. Paraffin crystal sizes were about 1200 µm^2^, much larger than the beam size. The higher-order reflections are clearly visible in Figure 3a after dosing with the pulsed electron beam compared to those with the continuous beam at the same total accumulated dose 600 e^−^·nm^−2^, suggesting the pulsed electron beam has a lower rate of radiation damage for the C_36_H_74_ orthorhombic crystal structure. To quantify this, the decrease in relative intensity of the {110} reflections (i.e., fading curve) was acquired for six different duty cycles at the same 5.2 GHz gate repetition rate. In Figure 3b, the fading curves are shown for the intensities of the {110} reflections (indexed four peaks on Figure 2a) from paraffin two-dimensional single crystals under continuous (100%, grey dots) and pulsed (22%, light-blue dots) beam mode. The critical dose was obtained by an exponential decay fit of the fading curves. Table 1 shows that the critical dose using 300 keV varies between 571 and 1124 e^−^·nm^−2^, for 100% and 10% duty cycles, respectively. This factor of 2 is comparable to the 1.8x dose rate improvement in the paraffin fading curve reported by University of Minnesota for 200 keV laser-generated single-electron packets ^14^.

**Table 1.** Critical doses (e^−^·nm^−2^) of C_36_H_74_ paraffin {110} reflections, and purple membrane (430) reflection, measured by intensity fading to 1/*e* for different duty cycles (%) at 5.2 GHz gate repetition rate from an exponential decay fit. Ratio 2 and 3 are the average of 200 and 300 keV results, respectively.

| *Sample* | *Duty Cycles ($D_f$)*<br>*@ 5.2 GHz*<br>*Electron*<br>*Energy* | *100%*<br>*(cont.*<br>*mode)* | *47%* | *32%* | *22%* | *14%* | *10%* |
| --- | --- | --- | --- | --- | --- | --- | --- |
| *C<sub>36</sub>H<sub>74</sub>*<br>*paraffin* | *300 keV* | 571.22 | 714.96 | 801.91 | 916.60 | 1046.55 | 1124.10 |
|  | *200 keV* | 441.12 | 542.55 | 602.10 | 683.41 | 778.96 | 830.98 |
|  | *Ratio 1*<br>*(300 keV/200 keV)* | 1.3±0.36 | 1.32±0.48 | 1.33±0.51 | 1.34±0.62 | 1.34±0.57 | 1.35 ±0.6 |
|  | *Ratio 2 [200 keV]*<br>*($D_f$ / cont.)* | 1 | 1.23±0.39 | 1.37±0.37 | 1.55±0.42 | 1.77±0.43 | 1.88±0.31 |
|  | *Ratio 3 [300 keV]*<br>*($D_f$ / cont.)* | 1 | 1.25±0.46 | 1.4±0.49 | 1.6±0.56 | 1.83±0.5 | 1.97±0.66 |
| *Purple*<br>*membrane* | *300 keV* | 232.01 | 298.04 | 346.13 | 391.73 | 450.45 | 489.00 |
|  | *200 keV* | 181.35 | 231.65 | 267.92 | 302.00 | 352.34 | 373.23 |
|  | *Ratio 1*<br>*(300 keV/200 keV)* | 1.28±0.15 | 1.29±0.14 | 1.29±0.21 | 1.3±0.24 | 1.28±0.26 | 1.31±0.18 |
|  | *Ratio 2 [200 keV]*<br>*($D_f$ / cont.)* | 1 | 1.28±0.09 | 1.48±0.12 | 1.67±0.16 | 1.94±0.14 | 2.06±0.1 |
|  | *Ratio 3 [300 keV]*<br>*($D_f$ / cont.)* | 1 | 1.29±0.21 | 1.49±0.24 | 1.69±0.23 | 1.94±0.28 | 2.11±0.23 |

**Figure 3.**
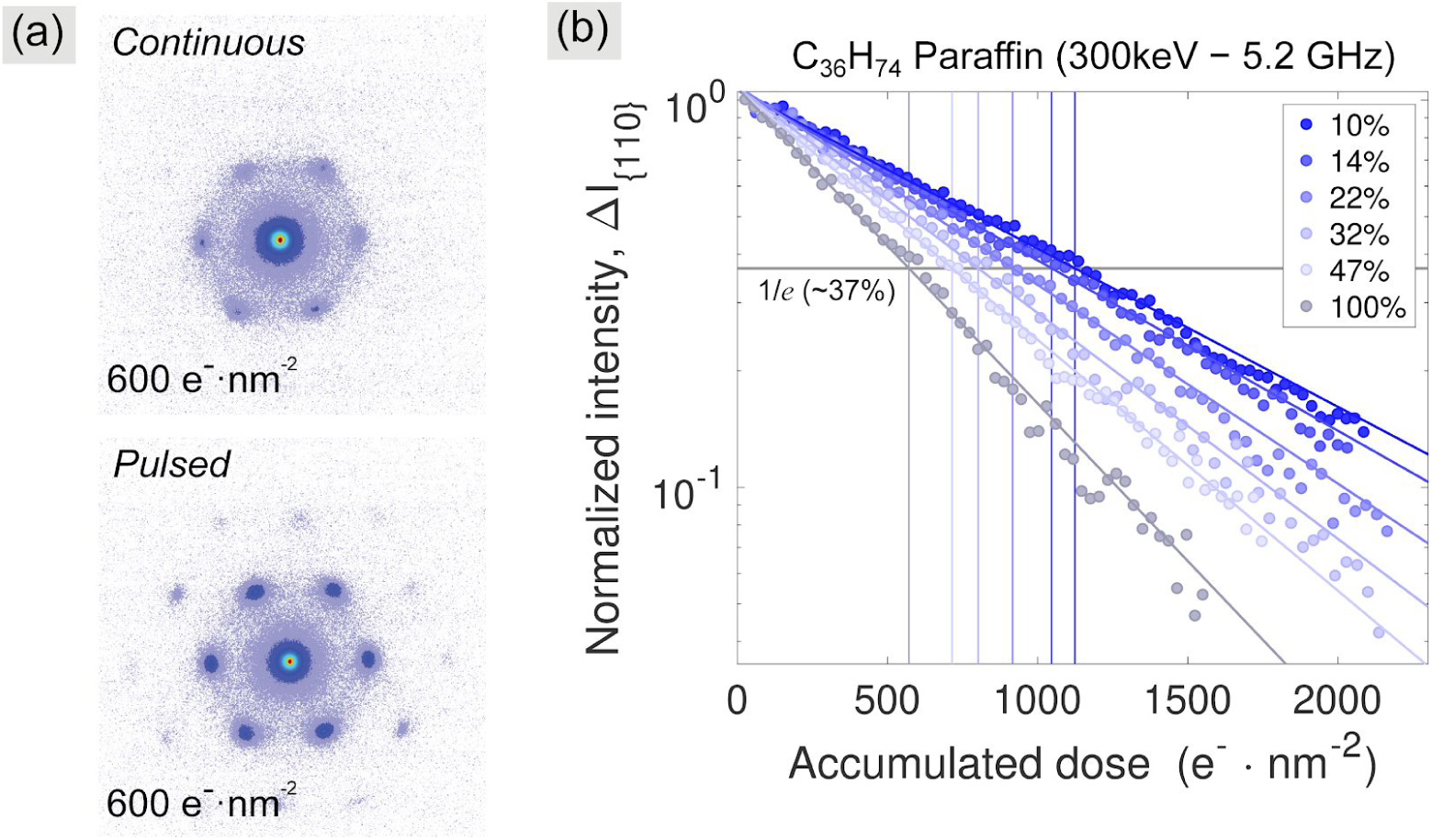
Mitigation of radiation damage at different duty factors of pulsed electrons. (a) Selected continuous- and pulsed-beam diffraction patterns of a paraffin crystal with the 110 family Bragg peaks highlighted. (b) Fading curves from the {110} reflections and exponential decay fits as a function of pulsed beam duty cycle factor at 5.2 GHz repetition rate for the same dose rate (5.11±0.01 e^−^·nm^−2^·s^−1^), grey guideline indicating critical dose (D_cri_).

The tunable GHz RF pulser achieved an effective time-averaged arrival rate of 73 MHz or higher for single-electron packets (see Table 2), 5 second acquisition time per each single diffraction image, and acquisition of a full pulsed-beam fading curve in a few minutes.

**Table 2.** Summarized experimental parameters.

| *Sample* | *Beam type* | *Freq. (GHz)* | *Duty cycle (%)* | *Avg. beam current (pA)* | *Average electrons / gate* | *Avg. arriv. freq. (MHz)* | *Beam area ($\mu\text{m}^2$)* | *Dose rate ($e^- \cdot \text{nm}^{-2} \cdot \text{s}^{-1}$)* | *Total acq. time (s)* |
| --- | --- | --- | --- | --- | --- | --- | --- | --- | --- |
| **Paraffin (C<sub>36</sub>H<sub>74</sub>)** | Contin. | N/A | N/A | 19.39 | N/A | N/A | 23.74 | 5.090 | 325 |
|  | Contin. | 5.2 | 100 | 100.13 | 0.120 | 624.0 | 122.72 | 5.093 | 270 |
|  | Pulsed | 5.2 | 47 | 46.78 | 0.056 | 291.2 | 56.98 | 5.125 | 320 |
|  | Pulsed | 5.2 | 32 | 32.53 | 0.039 | 202.8 | 40.02 | 5.074 | 295 |
|  | Pulsed | 5.2 | 22 | 22.14 | 0.027 | 140.4 | 27.05 | 5.109 | 300 |
|  | Pulsed | 5.2 | 14 | 14.36 | 0.017 | 88.4 | 17.39 | 5.154 | 325 |
|  | Pulsed | 5.2 | 10 | 10.38 | 0.014 | 72.8 | 12.65 | 5.122 | 350 |
| **Purple membrane** | Contin. | 5.2 | 100 | 100.04 | 0.120 | 624.0 | 298.65 | 2.091 | 320 |
|  | Pulsed | 5.2 | 47 | 47.12 | 0.056 | 291.2 | 136.48 | 2.155 | 320 |
|  | Pulsed | 5.2 | 32 | 32.62 | 0.039 | 202.8 | 100.07 | 2.035 | 340 |
|  | Pulsed | 5.2 | 22 | 21.97 | 0.027 | 140.4 | 65.31 | 2.100 | 340 |
|  | Pulsed | 5.2 | 14 | 14.11 | 0.017 | 88.4 | 41.25 | 2.135 | 365 |
|  | Pulsed | 5.2 | 10 | 10.13 | 0.014 | 72.8 | 30.60 | 2.066 | 365 |

Our method is continuously tunable in stroboscopic frequency, unlike RF resonator pulsers ^17^. For a given frequency, duty cycle is tunable (by varying RF power) as in the six different fading curves in Figure 3b. Exponential fits (Table 1) reveal duty cycle dependence of critical dose (D_cri_). Best results used short gates: 10% duty cycle at 5.2 GHz gate repetition rate doubled critical dose compared to continuous beam.

### Mitigation of purple membrane radiation damage by tunable pulsed beam

Radiation damage mitigation studies for purple membrane captured powder diffraction patterns at regular time intervals at room temperature (see Table 2 for 300 keV). Large illuminated areas containing many small (∼1µm^2^) purple membrane particles show powder diffraction rings characteristic of a circularly averaged hexagonal lattice with *p3* symmetry (a = b = 62.45 Å) ^25–28^. The time series were analyzed for the (430) reflection intensity (d-spacing = 8.827 Å) ^25^ versus accumulated electron dose. Intensities were normalized to their initial value and followed until fading completely.

Figure 4a shows that pulsed beam reduces radiation damage to purple membrane: (430) diffraction intensity is significantly stronger with pulsed compared to continuous beam after exposure to 230 e^−^·nm^−2^ accumulated dose. Rings are diffuse, due to a thick sample or soft phonon modes at room temperature. Figure 4b shows the (430) reflection fading curves for six duty cycles (gate widths) at 5.2 GHz gate repetition rate. Each exponential decay fit is taken at 1/e as the critical dose. As with paraffin, the critical dose (grey guideline) improves with smaller gate width (Table 1). Here, the 10% duty cycle improves the critical dose up to a factor of 2.1 at room temperature, similar to paraffin. Tuning the single-electron arrival rate is an effective damage mitigation strategy in both macromolecules.

**Figure 4.**
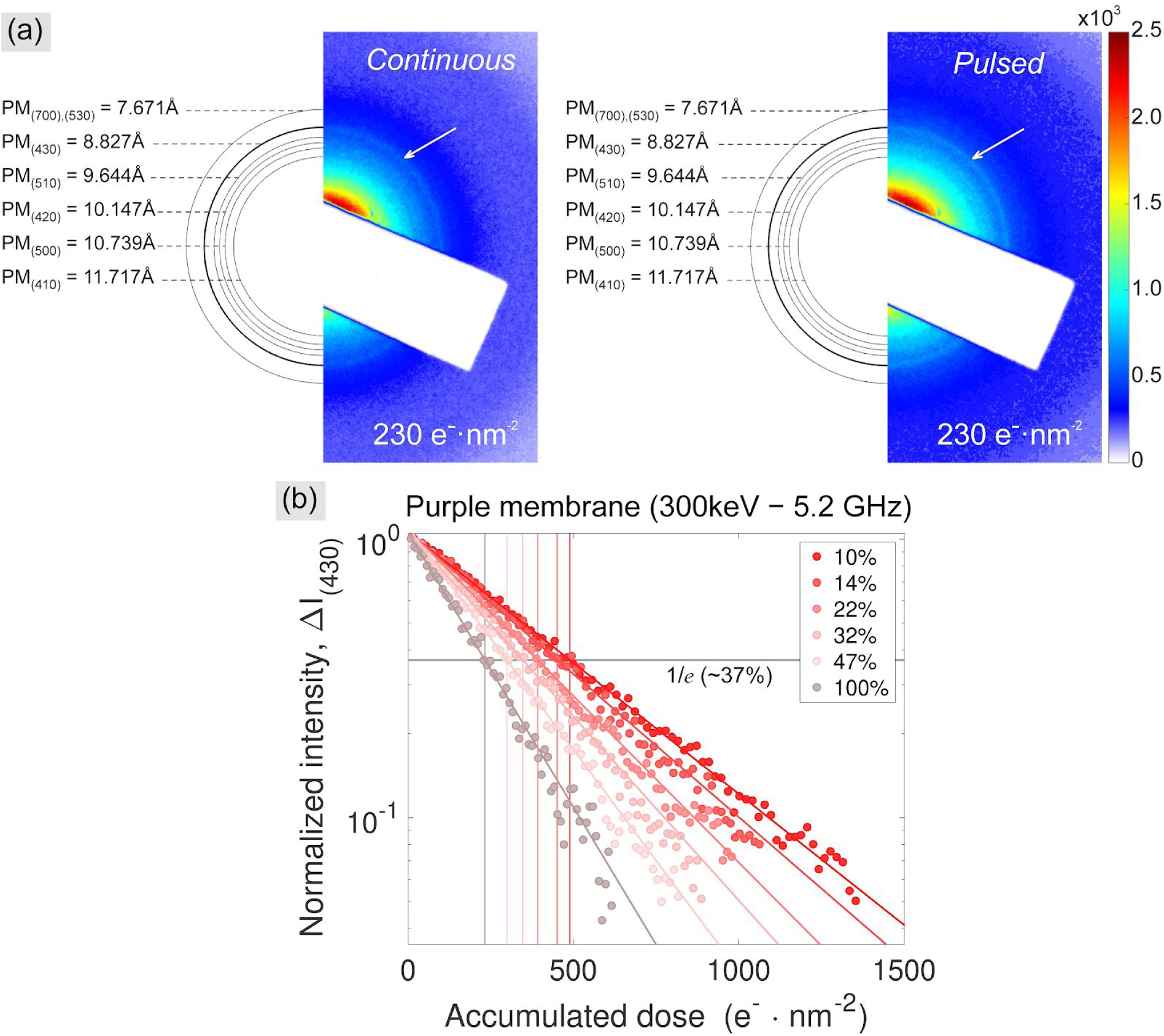
Comparison of the continuous random and pulsed random electron beam damage for the same dose rate (2.098±0.02 e^−^·nm^−2^·s^−1^). (a) Selected continuous- and pulsed-beam powder diffraction patterns of a purple membrane (PM) with the (430) reflection (marked arrows) at total accumulated dose 230 e^−^·nm^−2^. (b) Fading curves from the (430) reflection and non-linear exponential decay fits for each measured duty cycle factor from 10 to 100%.

### Energy dependence of radiation damage for both macromolecules using pulsed mode

Figure 5 compares the 10% duty factor pulsed mode fading curves of the relative intensity for both (a) organic C_36_H_74_ paraffin {110} reflections and (b) purple membrane (430) reflection at 300 keV and 200 keV. For comparison at two electron energies under pulsed mode, we performed the experiments at the NIST (300 keV) and BNL (200 keV) microscopes in exactly the same way using the same specimen on the same grid with the same pulsed mode conditions, 5.2 GHz repetition rate and 10% duty cycle factor. At each microscope, we simply moved onto a fresh single-crystal region on that same specimen. Further, the same holder modified with a Faraday cup was used to calibrate pulsed beam current in both instruments. We show in Table 1 the critical doses (D_cri_), defined as the total accumulated electron dose (e^−^·nm^−2^) at which the selected relative intensities is reduced to 1/*e* (∼37%) of their initial value. We estimate D_cri_ from the exponential decay fit as the dose at 1/*e* of the initial intensity.

**Figure 5.**
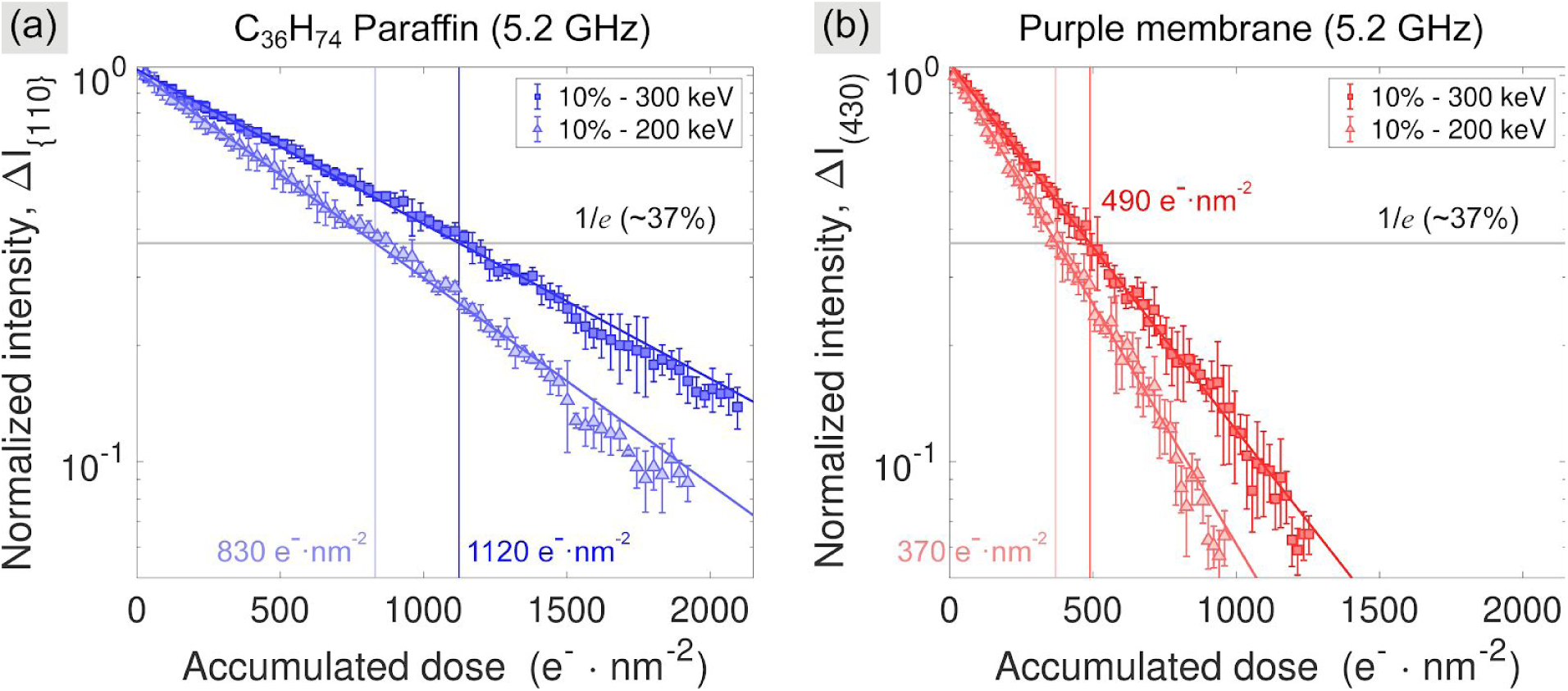
Comparison of electron radiation damage for 200 keV and 300 keV TEM in pulsed mode at 10% duty factor for both (a) organic C_36_H_74_ paraffin and (b) macromolecular purple membrane.

The calculated critical dose for C_36_H_74_ paraffin with 10% duty cycle pulsed mode is 1120 e^−^·nm^−2^ at 300 keV, and 830 e^−^·nm^−2^ at 200 keV. Averaging across identical trials, critical dose is 1.35±0.6 times higher at 300 keV than at 200 keV. Purple membrane has a similar energy ratio: 1.31±0.18. Continuous beam (Figure 2b) with paraffin had a 1.28±0.26 energy ratio. An energy-dependent slope is partially attributable to 1.13x higher inelastic scattering for carbon at 200 keV versus 300 keV ^13^, which for equal dose rates implies slower damage rates at 300 keV. Although purple membrane is more radiation-sensitive, its 2x damage mitigation with picosecond pulsed beam equals paraffin (Table 1), hinting at a common mitigation mechanism.

## Discussion

In this section we assess pulse-dependent critical dose at 5.2 GHz repetition rate for both 200 and 300 keV. We compare to other ultrafast TEM methods, discuss the meaning of tunable pulse structure from a single-electron perspective, and suggest future cryogenic applications.

Our key conclusion is that radiation damage mitigation using a GHz RF pulser is the new state of the art in time-structured low-dose TEM of organic macromolecules, effective for both C_36_H_74_ paraffin and purple membrane (bacteriorhodopsin) at room temperature and achieving record-fast stroboscopic image acquisition in single-electron mode. Full fading curves demonstrate remarkable, repeatable damage mitigation in macromolecules by pulsed vs continuous beam. Tuning the width of few-picosecond periodic gating of single electrons optimizes the low dose time structure. Critical dose doubles for both macromolecules in pulsed mode while holding dose rate constant, hinting at a common damage mechanism mitigated by picosecond pulse structure. This factor of ∼2 echoes the 1.8x dose rate improvement in paraffin reported by University of Minnesota for 200 keV laser-generated single-electron packets compared to continuous random beam with equal dose rate ^14^. But the GHz RF pulser can access the same time-structured single-electron regime - and the same damage mitigation - as the laser pulser at much higher repetition rates with orders of magnitude faster acquisition. Laser-based pulsed TEM is typically operated at <0.1GHz ^20^, limited in the cited study to 200 kHz single-electron pulse trains which took 22 hours to acquire only 2% of the paraffin 1/e fading curve ^14^. For comparable illuminated area on paraffin samples we achieved an effective time-averaged single-electron arrival rate of 73 MHz or higher (see Table 2), 5 second acquisition time per diffraction image, and a full 1/e time-series fading curve in a few minutes. Also, GHz RF pulsers, unlike laser-based systems, are compatible with field emission sources and have low energy spread, key for high resolution ultrafast stroboscopic TEM development. For a 10% duty cycle (10 pA beam), after converting intensity to electron counts by our Faraday cup calibration, we count 125 million electrons in just 2 s, of which 2.8 million are scattered away from the central spot. This is more than sufficient to obtain a diffraction pattern, and is repeated at 5 s intervals (for all data sets). Rapid acquisition in single-electron mode outperforms laser-based systems substantially, highlighting opportunities for ultra low-dose or pump-probe stroboscopic experiments using tunable picosecond pulsed electron beams.

Energy dependence is informed by our parallel experiments on 200 and 300 keV TEMs (each installation is fixed-energy due to phase matching the RF traveling wave to the electron velocity). The 300keV/200keV ratio of pulsed-beam critical dose at 10% duty factor for continuous beam paraffin, pulsed paraffin and pulsed purple membrane are 1.28±0.26, 1.35±0.6, and 1.31±0.18 respectively (Fig. 5). We note the inelastic scattering cross-section, which has an inverse proportionality with critical dose, has for carbon a 200keV/300keV ratio of 1.13 (and elastic, 1.27) ^13^. The slight mismatch between the energy ratio of inelastic scattering and the damage for a thin sample is within error bars, but could hint that nonlinear (even time-of-arrival dependent) effects may play a role and motivates an optimized picosecond time-structuring. Experiments with smaller error bars (likely requiring thinner samples) could reveal whether the critical dose energy dependence is identical for pulsed beam and continuous beam hence how it depends on inelastic scattering. Full results of the energy dependence of pulsed-mode critical dose for duty cycles from 10 − 100% are shown in Table 1. While critical dose for our bulk samples (drop-cast paraffin estimated thickness 18 nm by the Dorset method ^22^) is about 30% higher at 300 keV than 200 keV for both continuous or pulsed beam, the Peet-Henderson-Russo information coefficient is a better figure of merit than critical dose for selecting beam energy. Peet et al predicted an optimal information coefficient at 100 keV for thin (30 nm) single particle samples in vitreous ice; for thicker (100 nm) samples, 300 keV ^13^. The tunable GHz RF pulser can be designed to any TEM energy, for instance that with the highest information coefficient.

Here we analyze for the first time the correlation with critical dose of a wide range of average single-electron repetition rates (∼60 − 600 MHz). We are careful to distinguish the average single-electron repetition rate at these low beam currents, which is statistical, from the pulser gating repetition rate (5.2 GHz), which is fixed; the single-electron rate is lower than the gating rate because not all gates contain an electron. We model our beam with a Poisson time distribution in Figure 6a, showing the single-electron-per-pulse approximation holds at 10% duty cycle (leftmost point). Each point represents the modeled two-electron arrival probability within a single gate width versus calculated average arrival frequency for each experimental duty factor (Table 2). The single-electron-per-pulse approximation is good to 0.01% for gating at 10% duty factor. In Figure 6b we plot paraffin and purple membrane pulsed critical dose at 300 keV, normalized to continuous beam (from fits in Figures 3b and 4b), versus average arrival frequency of single electrons (Figure 6a). Each point is a different duty cycle. Improvements in critical dose show inverse dependence on average current, reaching 2x for <100 MHz, consistent with attributions of damage mitigation in the pulsed single-electron-packet regime to acoustic phonon dissipation ^17^ or local heating effects on radical diffusion ^14^. Sample-specific improvement suggests an optimum pulse structure varying by material: by tuning beam current, pulsed TEM frequency, and gate width (RF power), imaging and acquisition speed may be optimized.

**Figure 6.**
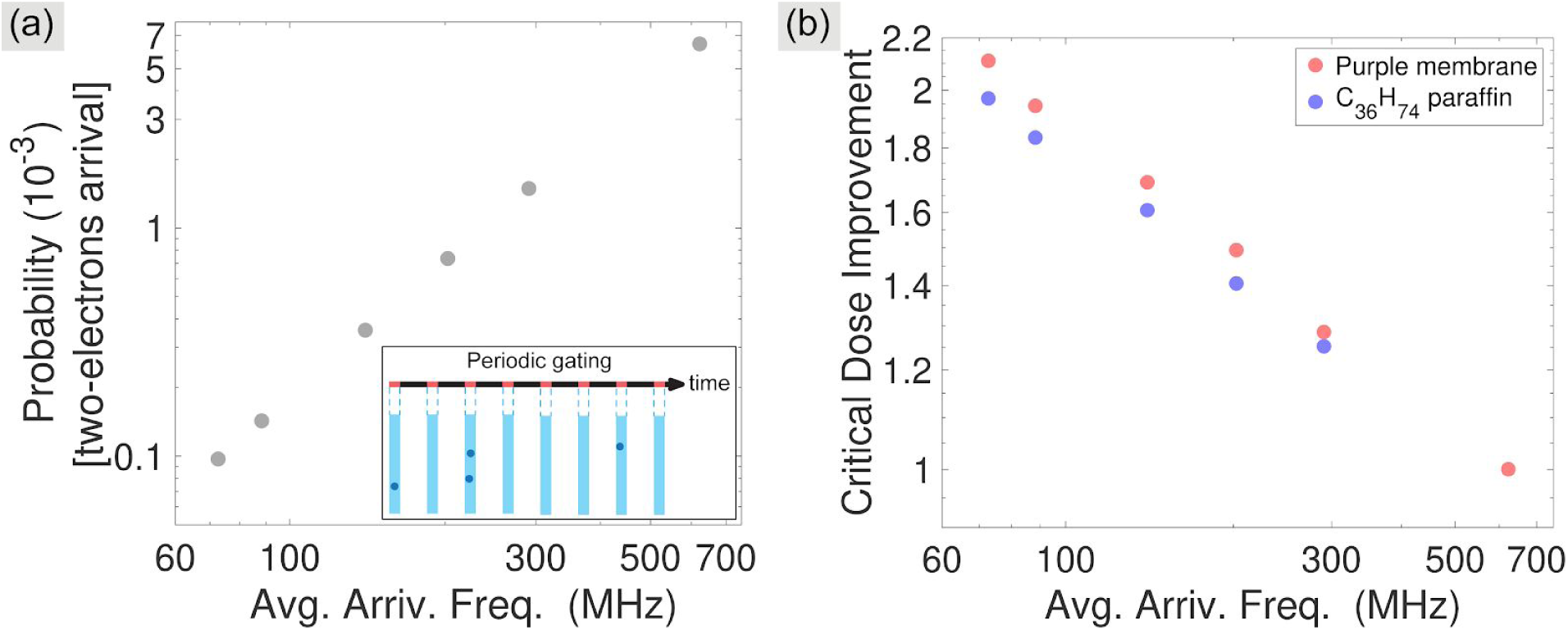
(a) Statistical model of electrons per gate at low current, as the Poisson probability of finding two random electrons in the same gate, 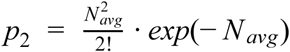, where *p*_*2*_ is the probability of arrival of 2 electrons given average electrons per window *N*_*avg*_. Probability is plotted versus average arrival frequency, which is calculated by multiplying the repetition rate with the average number of electrons per gate at a given current and duty cycle. (b) The measured critical dose improvement (Ratio 3 in Table 1) versus average arrival frequencies for both organic C_36_H_74_ paraffin and purple membrane using 300 keV stroboscopic TEM.

The statistical electron emission process (whether photo-, thermionic, or field emission) always leads to non-zero probabilities of multiple electron arrivals in short periods of time. Pulsing or gating restricts allowed arrival frequencies so single electrons are more likely.

We next examine whether reduction of double inelastic scattering - two scattering events within a few nanoseconds or less - correlates with damage mitigation. In addition to temporal separation, two scattering events will typically be spatially separated by a few tens of microns down to nanometers depending on the TEM magnification (spot size). We will not speculate on whether acoustic phonons or other phenomena can mediate a spatially-separated two-scatter effect, but simply discuss the frequency and timing of scattering events in our experiments alongside possible correlations with critical dose. In Figure 6, while the probability of multiple electrons arriving at the sample within one gating period correlates inversely with critical dose improvements, the magnitude of that probability is not correct for quantitative damage rate discussions. We estimate quantitatively as follows.

Double-electron arrival is a necessary but not sufficient condition for double scattering. For gates with two or more electrons arriving, usually a single electron will scatter or none at all; how many gates scatter two electrons? For our standard 2 seconds exposure per image at 5.2 GHz, 10.4 billion gates occur. For duty factor D_f_=10%, i.e. 10 pA, 0.01% of the gates have two electrons using Poisson statistics as in Figure 6, or 1.04 million gates. Scattering is a rough estimate because our samples are drop-cast with nonuniform thickness. Estimating local thickness by the Dorset method ^22^, we obtain, from Figure 2, a (200) to (110) reflection intensity ratio of 1.93 ± 0.17, yielding 18.4 ± 1.6 nm thickness for a representative sample. Taking a carbon atomic density for C_36_H_74_ paraffin (hexatriacontane) of *η*=34.2 nm^-3^, thickness t=18.4 nm, elastic scattering cross section given approximately by *σ*_e_=21/*β*^2^ pm^2^ where *β* is the relativistic velocity v/c, and inelastic scattering *σ*_i_∼*σ*_e_/4 in this energy range, we can estimate scatter probability of an electron 1-T=1-e^-t/*λ*^ for mean free path *λ*=1/[*η*(*σ*_e_+*σ*_i_)].^13^ Neglecting multiple scattering in this thin sample, at 200 keV we calculate 2.69% of electrons scatter; at 300 keV, 2.17%. Since the Poisson n-electron probability in one gate is 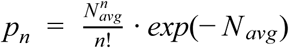 with 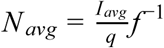 for electron charge q, average current I_avg_=10 pA (10% duty factor), and f=5.2 GHz, we obtain N_avg_=0.012, p_1_=0.010, and p_2_=0.00010. For one- and two-electron scattering within a single gate period, respectively, the event probability is *p*_1_ (1 − *T*) and *p*_2_(1 − *T*)^2^. The ratio of double to single interactions within one gate period is 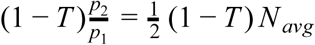 which is 1.3×10^−4^ at 10% duty factor and 300 keV, and is favorably smaller for thinner samples, shorter gates, or lower current. The ratio of double to single *inelastic* scattering interactions within one gate period (1/f=192 ps) is 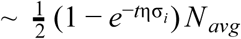 or 3.5×10^−5^ for 10% duty factor and 3.5×10^−4^ for continuous beam, an improvement of 10x. If two inelastic scattering events per gate (within 1/f = 192 ps) are 10x less likely at 10% duty factor, it does not explain the observed 2x critical dose. We suggest if double-inelastic-scatter is a mitigated damage mechanism here, the effective double-scatter time of importance in both paraffin and purple membrane is likely at least 5/f, or ∼1 ns.

The statistical chance of multiple electrons per pulse is important, but so too is the practical time required to obtain images. This reinforces a key advantage of our approach: the multi-electron probability can be kept as low as with laser-based systems, but our repetition rate is perhaps a thousand times higher while maintaining tunability. Thus images are acquired in a fraction of the time, TEM system stability and drift are much less critical, and the productivity of a machine retrofitted with a GHz pulser rather than a laser is much preferable when doing low-dose imaging.

All experiments presented here were performed at room temperature. But most single-particle imaging uses cryogenic electron microscopy (CryoEM) with samples suspended in low density amorphous (LDA) water. The beam-induced damage problem is severe for single-particle imaging because the critical dose for the sample is much smaller than that needed for high resolution images. Could pulsed-beam with a cryo-holder provide improvements to critical dose equaling or bettering the factor of two seen with pulsed beam at room temperature? If indeed a significant contributing factor to damage mitigation is phonon damping following single-electron interaction, or radical diffusion from local heating, this provides strong motivation for incorporating tunable GHz pulser technology into CryoEM. We specifically highlight at room temperature with pulsed beam our 10 nanosecond (100 MHz) average time between incident electrons, and our 50 microsecond (200 Hz) average time between electron scattering events, which doubled the critical dose versus non-pulsed beam with equal current density. On these timescales, faster damping or cooling of a cryogenic sample than is possible at room temperature might permit higher repetition rates, faster acquisition, and perhaps even higher critical dose. Material-dependent pulse optimization could be studied. In particular, the impact of pulsed beam irradiation on recrystallization of LDA water into ice, a perennial impediment to CryoEM imaging, would be of high interest. And for single-particle imaging, increased critical dose could enable higher resolution structural determination. As a first step towards these goals, cryo-holder measurements are planned and will be reported in a future work. Future cryo-compatible GHz pulsers are also planned at 100 keV, where the information/damage ratio is optimized for thin-sample single-particle CryoEM ^13^ and where our doubling of the damage threshold, should it hold or improve at that energy and temperature, could represent a new state of the art.

## Methods

### Sample preparation

Two model biological macromolecular specimens were used in this study. The organic material is C_36_H_74_ paraffin, which forms thin, orthorhombic single crystals oriented along the [001] crystallographic direction upon drop-casting from solution onto a substrate. The paraffin solution was prepared by dissolving C_36_H_74_ paraffin powder (98% purity *n*-hexatriacontane, Sigma-Aldrich) in *n*-hexane solvent (95% purity, Consolidated Chemical & Solvents LLC). Samples for use in TEM were prepared by pipetting a droplet of the C_36_H_74_ paraffin solution onto carbon-coated copper (Cu) 300 mesh TEM grids (Lacey Carbon 300 Mesh Copper Grids, 3mm, SPI Supplies) followed by solvent evaporation in air.

Purple membrane was isolated from H. salinarum strain S9 by the standard method ^29^ and diluted to 3 mg/ml in 0.1 M potassium phosphate buffer at pH 5.2 ^30^. All the samples were provided by Drs. Joseph Mckeown and Megan Shelby (Lawrence Livermore National Laboratory). The purple membrane solution was washed by pelleting via centrifugation at 12000 rpm for 1.5 hours until the supernatant was clear and then resuspended in water. The sample was deposited onto a carbon-coated Cu mesh (Holey Carbon 300 Mesh Copper Grids, 3mm, SPI Supplies) by dropping 2 μl of 2 mg/ml in 2% glucose in water onto the grid and allowing the solution to fully air dry without blotting.

### GHz pulser operation and electron diffraction pattern acquisition

Electron crystallography experiments were performed at two stroboscopic TEM facilities to ensure repeatability: a modified 300 keV LaB_6_-based thermionic TEM (JEM-3010, JEOL, with a Euclid STROBO-300) and a modified 200 keV Schottky-emission TEM (JEM-2010F, JEOL, with a Euclid STROBO-200). 5.2 GHz duty cycle was calibrated for each machine using a Faraday cup and a 6517B Keithley picoammeter. By using a GHz radio-frequency (RF) microwave source to precisely control the electron chopping process, electron packets as short as 20 ps are delivered at tunable GHz repetition rates; here the electron beam current was reduced to tens of pA; single electron arrival rate is preferred terminology vs pulse width (Table 2). To be clear, we are not observing a simple low-dose single-electron effect: picosecond gating of single electron arrival is what doubles critical dose, as shown in Table 2 for constant dose rate (flux) comparing pulsed and continuous beams.

All diffraction patterns at 300 keV were recorded by Gatan Orius SC1000 charge-coupled device (CCD) camera at room temperature using an indicated camera length of 300 mm, and at 200 keV by a Gatan UltraScan 1000XP. The electron beam current in the single-electron-packet regime was between 10 and 100 pA and the illuminated area approximately 4−19.5 µm in diameter (measured accurately using a cross grating replica for calibration of magnification), with illuminated area reduced in proportion to average current through the dose series to keep flux constant (Table 2). The series of diffraction images was automatically saved by the ‘*acquire series*’ data-acquisition function of the Gatan DigitalMicrograph operation system. In this study, we used 5 s acquisition time per each image, containing ∼ 3 s CCD read-out time plus 2 s detector exposure time. Gatan DigitalMicrograph and MATLAB software were used for image capture and analysis, respectively.

For C_36_H_74_ paraffin, a series of 54−70 images × 5 s acquisition time was obtained at a flux of 5.11 e^−^·nm^−2^·s^−1^. Beam-induced damage was studied by locating a fresh area of paraffin crystal in bright field mode, reducing spot size in proportion to average current to keep flux constant, followed by capture of diffraction patterns at set time intervals until diffraction peaks had faded ^14,31^. The series of diffraction patterns were analyzed by measuring the intensity of pairs of Bragg peaks (typically the {110} reflections contained four peaks: marked indices on Figure 2a) at each time interval and relating that to the accumulated electron dose up to that time. Intensities were averaged and normalized against the initial highest intensity. The accumulated electron dose (D_accum_) is determined as 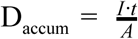, where *I* is beam current (e^−^·s^−1^), t is the acquisition time (s), and A is the illuminated beam area (nm^−2^).

For purple membrane, a series of 64−73 images × 5 s acquisition time at a flux of 2.098 e^−^·nm^−2^·s^−1^ was taken. Electron diffraction rings of purple membrane were analyzed by a radial averaging method ^13,30,32^, then the average intensities in one resolution zone (centered at 8.827 Å) were normalized to their initial intensity and followed as a function of accumulated electron dose. All the experimental parameters discussed in this report are summarized in Table 2.

## Data Availability

All the data that support the findings of this study are available within the paper, or from the corresponding author upon reasonable request.

## Acknowledgements

This work was supported by the SBIR program under NIH project number 1R43GM133267-01. The research at BNL was supported by the U.S. Department of Energy, Office of Basic Energy Sciences, Division of Materials Science and Engineering, under Contract No. DE-SC0012704. The NIST microscope was commissioned under Award No. SB1341-16-CN-0035. The authors would like to thank Dr. Michael B. Katz of NIST for his kind support and Dr. Myung-Geun Han for assistance with 200 keV JEM-2010F operation at BNL, as well as Dr. Joseph Mckeown and Dr. Megan Shelby of Lawrence Livermore National Laboratory for providing purple membrane samples and for useful discussions.

## Notes

### Competing Interest Statement

The authors have declared no competing interest.

